# Severely ill COVID-19 patients display augmented functional properties in SARS-CoV-2-reactive CD8_+_ T cells

**DOI:** 10.1101/2020.07.09.194027

**Authors:** Anthony Kusnadi, Ciro Ramírez-Suástegui, Vicente Fajardo, Serena J Chee, Benjamin J Meckiff, Hayley Simon, Emanuela Pelosi, Grégory Seumois, Ferhat Ay, Pandurangan Vijayanand, Christian H Ottensmeier

**Author notes:** These authors jointly contributed to the work (Anthony Kusnadi, Ciro Ramírez-Suástegui, Vicente Fajardo, Serena J Chee). These authors jointly directed the work (Pandurangan Vijayanand, Christian H Ottensmeier). Correspondence: P.V. or C.H.O.

## Abstract

The molecular properties of CD8_+_ T cells that respond to SARS-CoV-2 infection are not fully known. Here, we report on the single-cell transcriptomes of >80,000 virus-reactive CD8_+_ T cells from 39 COVID-19 patients and 10 healthy subjects. COVID-19 patients segregated into two groups based on whether the dominant CD8_+_ T cell response to SARS-CoV-2 was ‘exhausted’ or not. SARS-CoV-2-reactive cells in the exhausted subset were increased in frequency and displayed lesser cytotoxicity and inflammatory features in COVID-19 patients with mild compared to severe illness. In contrast, SARS-CoV-2-reactive cells in the non-exhausted subsets from patients with severe disease showed enrichment of transcripts linked to co-stimulation, pro-survival NF-κB signaling, and anti-apoptotic pathways, suggesting the generation of robust CD8_+_ T cell memory responses in patients with severe COVID-19 illness. CD8_+_ T cells reactive to influenza and respiratory syncytial virus from healthy subjects displayed polyfunctional features. Cells with such features were mostly absent in SARS-CoV-2 responsive cells from both COVID-19 patients and healthy controls non-exposed to SARS-CoV-2. Overall, our single-cell analysis revealed substantial diversity in the nature of CD8_+_ T cells responding to SARS-CoV-2.

## INTRODUCTION

Coronavirus infections with SARS-CoV-2 have created a global crisis; a large international effort is underway to develop treatments and vaccines to reduce the severity of disease and to provide protective immunity. To inform this effort, a detailed understanding of anti-viral immune responses is required. CD8_+_ T cell responses are thought to be critical for control of viral infections^1–4^, but to date, our understanding of anti-viral CD8_+_ T cell responses, specifically against Coronaviridae during infection and in the memory phase, is limited. Recently, studies have begun to improve our knowledge about CD8_+_ T cells responsive to SARS-CoV-2^5–14^, but the molecular features that associate with poor clinical outcomes or differentiate them from other virus-reactive CD8_+_ T cells remain incompletely understood. We report here on the data generated by single-cell RNA sequencing of virus-reactive CD8_+_ T cells from COVID-19 patients with varying severity of disease. We benchmark these data against the transcriptomes from CD8_+_ T cells from healthy donors, who have memory responses to other respiratory viruses.

## RESULTS

### Evaluation of virus-reactive CD8_+_ T cells

From 39 subjects with confirmed SARS-CoV-2 infection (17 patients with relatively milder disease not requiring hospitalization, 13 hospitalized patients and 9 additional patients requiring intensive care unit (ICU) support (**Fig. 1a** and **Extended Data Table 1**)), we isolated virus-reactive CD8_+_ memory T cells using a modified Antigen-Reactive T cell Enrichment (ARTE) assay^15–17^. Peripheral blood mononuclear cells (PBMC) were first stimulated for 24 hours with peptide pools specific to SARS-CoV-2^7,8^ (online Methods). Responding cells were then isolated based on the expression of the cell surface activation markers CD137 and CD69 (**Fig. 1a,b** and **Extended Data Fig. 1a**)^5,8,9^. We observed that the numbers of SARS-CoV-2-reactive CD8_+_ memory T cells were significantly increased in patients with severe COVID-19 illness who required hospitalization compared to those with milder illness not requiring hospitalization (**Fig. 1b**). A large fraction of SARS-CoV-2-reactive CD8_+_ T cells co-expressed CD279 (PD-1), CD38, and HLA-DR, which are markers linked to T cell activation and exhaustion^6,18,19^ (**Fig. 1c, Extended Data Fig. 1b** and **Extended Data Table 2**). Recent studies in patients with COVID-19 illness have reported that circulating CD8_+_ T cells express activation markers CD137 and CD69, likely activated by SARS-CoV-2 infection *in vivo*^5,9^; these cells are also captured and contribute to an unbiased evaluation of virus-reactive T cells from patients with recent COVID-19 illness.

**Figure 1.**
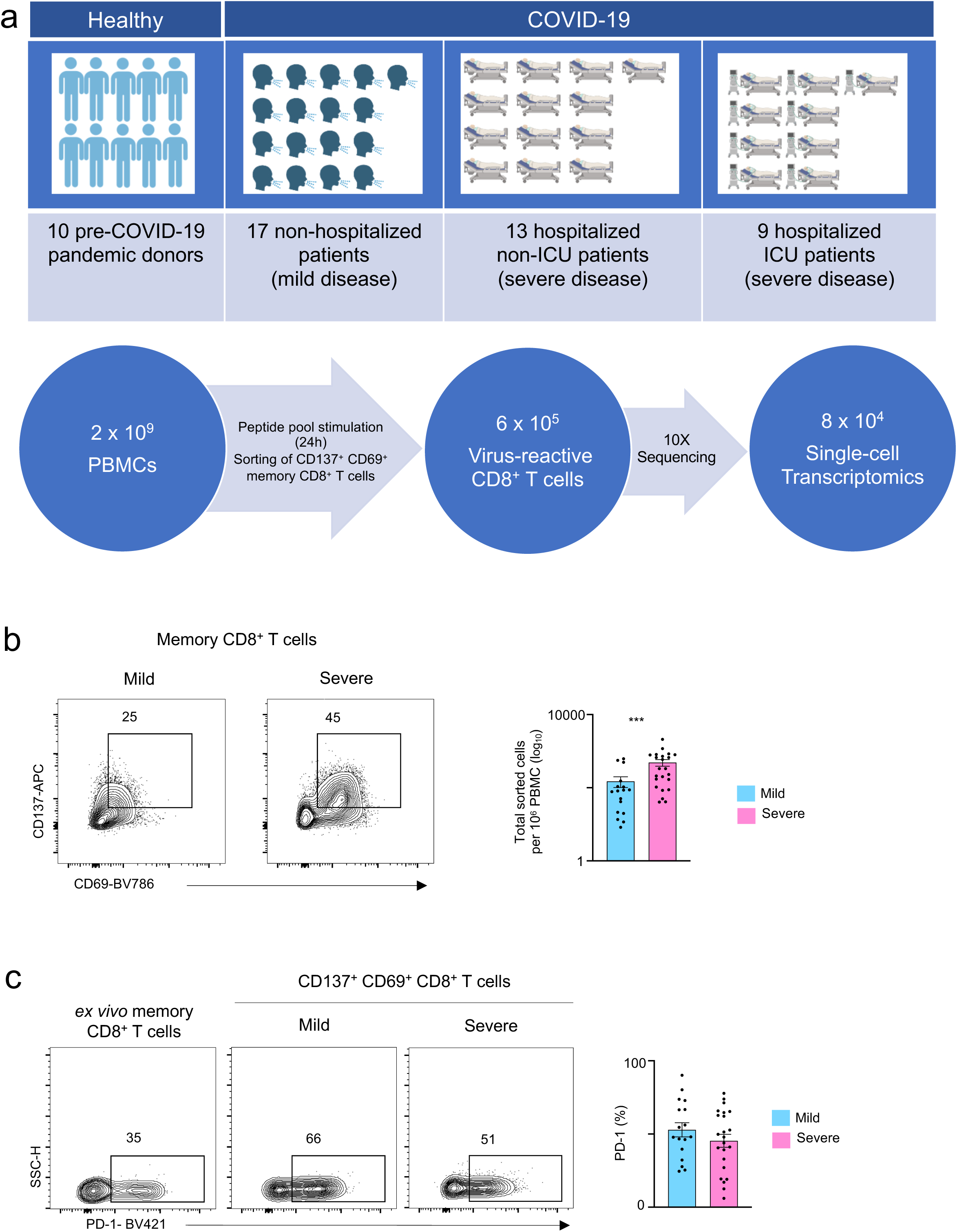
CD8_+_ T cell responses in COVID-19 illness. **a**, Study design overview. **b**, Representative FACS plots displaying surface staining of CD137 and CD69 in post-enriched CD8_+_ memory T cells, stimulated for 24 hours with SARS-CoV-2 peptide pools, from COVID-19 patients with mild and severe illness (left), and summary of the number of cells sorted per million PBMC (right). **c**, Representative FACS plots (left) showing surface expression of PD-1 in CD8_+_ memory T cells *ex vivo* (without *in vitro* stimulation) and in CD137_+_CD69_+_ CD8_+_ memory T cells following stimulation, post-enrichment (CD137-based) and corresponding summary plots (right) showing proportion of PD-1 expressing cells in each study subject (*P* = 0.26, unpaired t-test); Data are displayed as mean +/- S.E.M (N=39). \*\*\**P*<0.001 by Mann-Whitney test (**b**).

To study the properties of SARS-CoV-2-reactive CD8_+_ T cells in healthy non-exposed individuals^5,20,21^, we isolated CD8_+_ T cells responding to SARS-CoV-2 peptide pools from healthy subjects, who provided blood samples pre-COVID-19 pandemic (**Fig. 1a** and **Extended Data Table 1**). To contextualize our data and to define shared or distinguishing properties of CD8_+_ T cells reactive with other common non-corona respiratory viruses, we stimulated PBMC from healthy subjects with peptide pools specific to respiratory syncytial virus (RSV) and influenza A (FLU) and isolated responding cells (**Fig. 1a**). In total, from 49 subjects, we sorted and analyzed the single transcriptome and T cell receptor (TCR) sequence of > 87,000 virus-reactive CD8_+_ T cells with good quality metrics (**Fig. 1a**, **Extended Data Fig. 1c,d** and **Extended Data Table 3**).

### Virus-reactive CD8_+_ T cells show transcriptomic heterogeneity

Our unbiased single-cell transcriptomic analysis of all the virus-reactive CD8_+_ T cells revealed 8 distinct clusters (**Fig. 2a-c** and **Extended Data Table 4**), indicating that CD8_+_ memory T cells can activate a wide range of transcriptional programs in response to different viral infections^22^. Recent reports from COVID-19 patients have suggested the presence of exhaustion-related markers in global CD8_+_ T cell populations. To examine, whether such ‘exhausted’ cells were present in our dataset, we first generated a consensus list of ‘exhaustion’ signature genes from 9 studies^23–31^ that reported exhaustion features in T cells analyzed from patients with infection or cancer and in mouse models of viral infection (**Extended Data Table 5**). Gene set enrichment analysis (GSEA) of individual clusters showed significant positive enrichment of exhaustion signature genes only in cluster 1 cells (**Fig. 2d-f** and **Extended Data Fig. 2a**). This cluster (cluster 1) was also highly enriched for genes linked to type I interferon signaling (**Fig. 2d-f** and **Extended Data Fig. 2a**), highlighting the association between the interferon signaling and exhaustion program that has been previously described in the context of murine LCMV infection^32–34^. Because CD4_+_ T cell-mediated help is required for the generation of robust memory CD8_+_ T cells^35^, we assessed if cells in the ‘exhausted’ cluster 1 displayed ‘unhelped’ features that are linked to the lack of CD4_+_ T cell ‘help’^35^. As expected, the exhausted’ cluster 1 was also significantly enriched for transcripts linked to ‘unhelped’ CD8_+_ T cells (**Fig. 2d,e**). Despite displaying ‘exhausted’ and ‘unhelped’ features, intriguingly, cells in cluster 1 showed significant positive enrichment of cytotoxicity signature genes and higher expression levels of cytotoxicity-associated transcripts such as *GZMB* and *GZMA* relative to other clusters (**Fig. 2d-f** and **Extended Data Fig. 2a,b**), which suggested potential heterogeneity within this exhausted subset.

**Figure 2.**
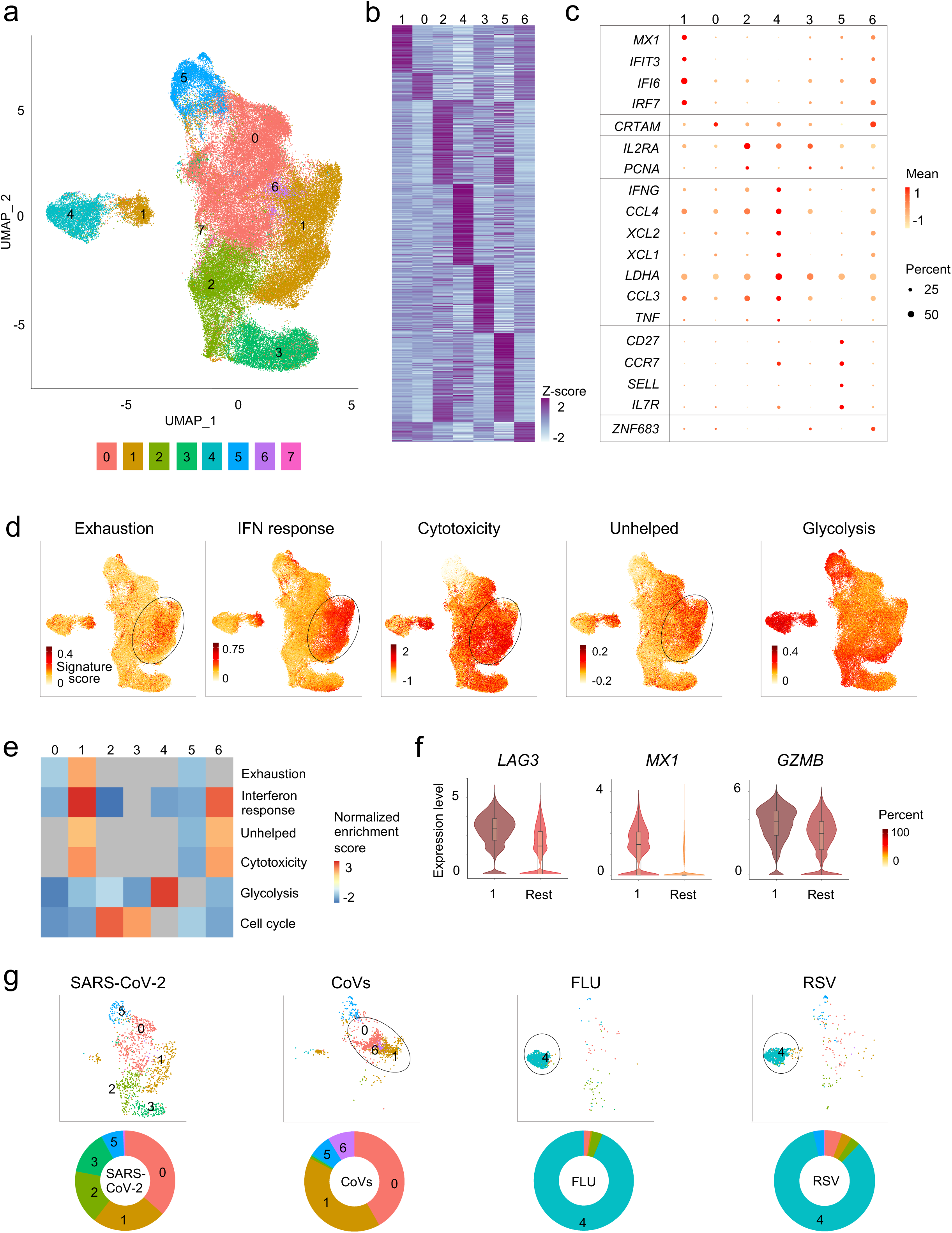
Virus-reactive CD8_+_ T cells show transcriptomic heterogeneity. **a**, Uniform manifold approximation and projection (UMAP) analysis that displays single-cell transcriptomic landscape of sorted CD137_+_CD69_+_ CD8_+_ memory T cells following 24 hours stimulation with virus-specific peptide pools. Seurat-based clustering of single cells colored based on cluster type. **b**, Heatmap showing expression of the most significantly enriched transcripts in clusters 0-6 (see **Extended Data Table 4**, Seurat marker gene analysis – comparison of a cluster of interest versus all other cells). Shown are a subset of the top 200 transcripts that have an adjusted *P* < 0.05, log_2_ fold change > 0.25, and >10% difference in the percentage of cells expressing the differentially expressed transcript between two groups compared. **c**, Graph showing average expression and percent expression of selected marker transcripts in each cluster; cells in cluster 7 that comprise <1% of all cells are not shown (**b,c**). **d**, UMAP is illustrating exhaustion, interferon (IFN) response, cytotoxicity, ‘unhelped’, and glycolysis signature scores for each cell. **e**, Gene Set Enrichment Analysis (GSEA) for the indicated signature genes comparing each cluster with the rest of the cells. Heatmap shows summary of the normalized enrichment scores for each cluster. Gray color indicates that the signature does not reach statistical significance (*P* >0.05) in a given cluster. **f**, Violin plots showing expression of representative exhaustion, IFN response, cytotoxicity marker transcripts (*LAG3, MX1, GZMB*, respectively) in cluster 1 compared to an aggregation of remaining cells (Rest). The color indicates percentage of cells expressing indicated transcript. **g**, UMAPs are depicting CD8_+_ memory T cells for individual virus-specific pool stimulation conditions (top panel). Each group of virus-reactive cells was randomly downsampled to ensure equal representation, corresponding pie charts, displaying proportions of virus-reactive cells in individual clusters (bottom panel).

CD8_+_ T cells reactive to specific viruses made strikingly different contributions to individual clusters (**Fig. 2g** and **Extended Data Table 3**). Clusters 0,1,2, and 3 were overrepresented by SARS-CoV-2-reactive CD8_+_ T cells from COVID-19 patients, and are likely to reflect memory or effector cells generated in the context of a recent infection. In contrast, the vast majority (>80%) of FLU-reactive and RSV-reactive CD8_+_ T cells were present in cluster 4, where SARS-CoV-2-reactive cells were underrepresented (<2%) (**Fig. 2g**). Cells in cluster 4 expressed higher levels of transcripts encoding for cytokines such as IFN-γ, TNFα, CCL3, CCL4, XCL1, and XCL2 (**Fig. 2c**, **Extended Data Fig. 2c**, and **Extended Data Table 4**), resembled polyfunctional CD8_+_ T cells that have been linked to protective immunity against a range of viral infections^36–38 39^. Besides, this cluster displayed positive enrichment and the highest score for genes in the aerobic glycolysis pathway, which is linked to better effector function through multiple mechanisms that are independent of metabolism itself^40,41^ (**Fig. 2d,e** and **Extended Data Fig. 2a**).

SARS-CoV-2-reactive CD8_+_ T cells from healthy non-exposed subjects, presumed to be human coronavirus (CoV)-reactive cells that cross-react with SARS-CoV-2^21^, were mainly present in clusters 0,1, 5 and 6 (**Fig. 2g** and **Extended Data Table 3**). The cells in cluster 6 were highly enriched for the expression of transcripts encoding ZNF683 (**Extended Data Fig. 2d**), also known as HOBIT, a transcription factor that plays a pivotal role in the development of tissue-resident memory (T_RM_) cells^42^. Notably, the vast majority of cells in cluster 6 were from SARS-CoV-2-reactive cells of healthy nonexposed subjects (**Fig. 2g**); in contrast, cells from COVID-19 patients were mostly absent in this cluster. Overall, our data revealed substantial heterogeneity in the nature of CD8_+_ T cell subsets generated in response to different viral infections.

### Exhausted SARS-CoV-2-reactive CD8_+_ T cells are increased in mild COVID-19 illness

By linking single-cell transcriptome with T cell receptor (TCR) sequence data of the same cells, we observed extensive clonal expansion as well as clonal sharing of TCRs between the different SARS-CoV-2-reactive subsets in COVID-19 patients (**Fig. 3a,b** and **Extended Data Table 6,7**). Single-cell trajectory analysis showed that cells in cluster 0,1 and 2 were interconnected rather than following a unidirectional path (**Fig. 3c**). Together, our data support both diversity as well as plasticity in the nature of memory CD8_+_ T cell responses to SARS-CoV-2 infection. However, the dominant memory subsets varied substantially across COVID-19 patients with cells in some subsets represented only by a few patients (**Fig. 3d** and **Extended Data Table 3**). For example, over 85% of SARS-CoV-2-reactive CD8_+_ T cells in patient 8 were just from cluster 3 (identified by * in **Fig. 3d**), indicating a lack of plasticity in CD8_+_ T cell responses in this person.

**Figure 3.**
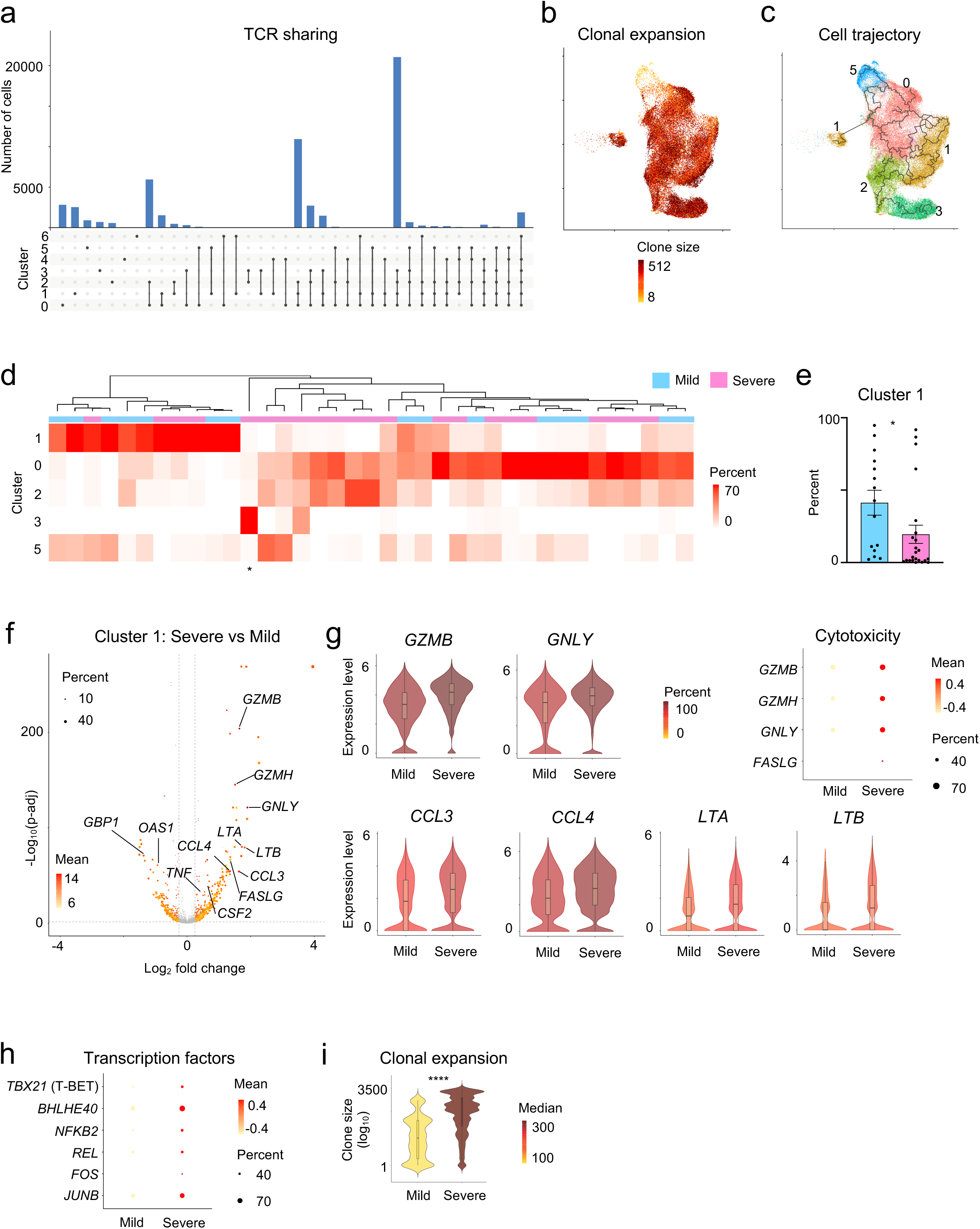
Exhausted SARS-CoV-2-reactive CD8_+_ T cells are increased in mild COVID-19 illness. **a**, Single-cell TCR sequence analysis of SARS-CoV-2-reactive cells showing the sharing of TCRs between cells from individual clusters (rows, connected by lines). Bars (top) indicate the number of cells intersecting indicated clusters (columns). **b**, UMAP is showing the clone size of SARS-CoV-2-reactive cells from COVID-19 patients. **c**, Single-cell trajectory analysis showing the relationship between cells in different clusters (line). **d**, Unsupervised clustering of all COVID-19 patients (mild and severe illness) based on the proportion of SARS-CoV-2-reactive CD8_+_ T cells present in each cluster per patient. The symbol * below represents patient 8. Clusters 4, 6, and 7 that had a very low frequency of cells in COVID-19 patients (<1% cells per cluster in total) are not shown here, full details provided in **Extended Data Table 3**. **e**, Bar chart comparing the proportion of cells in cluster 1 from COVID-19 patients with mild and severe illness. Data are displayed as mean +/- S.E.M (N=37, 2 subjects without hashtag data were not included for donor-specific analysis). **f**, Volcano plot showing genes differentially expressed (adjusted *P* < 0.05, mean CPM >0, log_2_ fold change >0.25) in cluster 1 cells between COVID-19 patients with severe and mild disease. **g**, Violin, and dot plots comparing the expression of indicated transcripts in cluster 1 cells between COVID-19 patients with mild and severe disease. **h**, Plot displaying the expression of several key transcription factors in cluster 1 cells from COVID-19 patients with severe and mild illness. **i**, Violin plots showing the degree of CD8_+_ T cell-clonal expansion in cluster 1 cells between COVID-19 patients with mild and severe disease. \**P*<0.05, \*\*\*\**P*<0.0001 by Mann-Whitney tests (**e**, **i**).

We next examined how SARS-CoV-2-reactive cell subsets varied between patients. COVID-19 patients broadly clustered into two groups based on whether the majority (>50%) of their CD8_+_ memory T cell responses to SARS-CoV-2 were either from cluster 1 or cluster 0 (**Fig. 3d**). Cluster 1 represented ‘exhausted’ cells with significant positive enrichment of both exhaustion and interferon signatures, whereas cluster 0 cells showed significant negative enrichment of these signatures, *i.e*., these cells were not ‘exhausted’ (**Fig. 2d,e**). Our analysis suggested that a sub-group of COVID-19 patients (30%) mounted a predominantly ‘exhausted’ CD8_+_ memory T cell response to SARS-CoV-2. The magnitude of this ‘exhausted’ response did not show a strong correlation to the time interval between onset of illness and sample collection (**Extended Data Fig. 3a**). Importantly, patients with milder disease had a significantly higher frequency of cells in the ‘exhausted’ cluster (cluster 1) when compared to those with severe disease (mean 41% versus 20%, **Fig. 3e**). Further, patients with severe disease when compared to those with mild disease showed significant enrichment of cytotoxicity signature genes and depletion of interferon signature genes in cluster 1 (**Extended Data Fig. 3b,c**), suggesting both quantitative and qualitative differences in cells in the ‘exhausted’ cluster based on disease severity. In support of qualitative differences, single-cell differential gene expression analysis showed that cells in the ‘exhaustion’ cluster (cluster 1) from severe COVID-19 patients expressed significantly higher levels of transcripts encoding for cytotoxicity-associated molecules (granzyme B, granzyme H, granulysin, Fas ligand^5,43^) and pro-inflammatory cytokines (CCL3, CCL4, CSF-2, TNF, LTA and LTB)^5,9,44^ (**Fig. 3f,g**, and **Extended Data Fig. 3d**). Transcripts encoding for several transcription factors that support cytokine production, inflammation and persistence (T-BET, BHLHE40, NFKB2, REL, FOS, JUNB)^45–50^ (**Fig. 3h,i** and **Extended Data Table 8**) were also expressed at significantly higher levels in cluster 1 cells from COVID-19 patients with severe illness. TCR sequence analysis of cells in the ‘exhausted’ cluster 1 revealed greater clonal expansion in patients with severe compared to mild illness, suggesting greater proliferative capacity and/or persistence of cells from severe COVID-19 patients (**Fig. 3i**). Given the importance of exhaustion programs in preventing excessive host tissue damage in viral infections^51,52^, we speculate that the failure to imprint an ‘exhaustion’ program that restrains T cell effector function may reflect a failure to limit exaggerated CD8_+_ T cell effector function, and thereby contribute to disease pathogenesis in some patients with severe COVID-19 illness.

### Pro-survival features in SARS-CoV-2-reactive CD8_+_ T cells from patients with severe COVID-19

SARS-CoV-2-reactive cells from COVID-19 patients, who did not mount a pre-dominant ‘exhausted’ response, were present mainly in clusters 0 and 2, the ‘non-exhausted’ subsets (**Fig. 2d, 3d**). These ‘non-exhausted’ subsets displayed cytotoxicity signature scores comparable to other clusters (**Fig. 2d**). Furthermore, cluster 2 was enriched for cell cycle signature, indicative of a higher proportion of proliferating cells in this cluster (**Fig. 2c, e**). We found no significant difference in the proportions of cells in cluster 0 and 2 between patients with severe and mild COVID-19 illness (**Extended Data Fig. 4a**).

However, single-cell differential gene expression analysis in clusters 0, as well as cluster 2, revealed significant transcriptional differences between COVID-19 patients with mild and severe illness (**Fig. 4a** and **Extended Data Table 8**). Ingenuity pathway analysis of transcripts with increased expression in cluster 0 from COVID-19 patients with severe relative to mild illness showed significant enrichment of transcripts in multiple co-stimulation pathways (OX40, CD27, CD28, 4-1BB, CD40), the NF-κB and apoptosis signaling pathways (**Fig. 4b-d**, **Extended Data Fig. 4b-d** and **Extended Data Table 9**). Co-stimulation is required for the robust activation and generation of memory T cell responses^53^, and activation of the NFκB pathway is important for T cell IL-2 production, proliferation, survival, cytokine production, and effector function^46^. IL2 receptor alpha and STAT5 encoding transcripts were also increased in severe compared to mild disease, indicating a greater potential for these cells to receive the pro-survival IL-2 signals^54^ (**Fig. 4a,c**). Other transcripts with increased expression encoded for transcription factors involved in cell fitness and cytokine production (BHLHE40)^45^, effector differentiation (BLIMP1)^55^ and prevention of exhaustion (JUN)^49^, and for CRTAM that has previously been shown to be important for generating effective cytotoxic T cells and viral clearance in mouse models^56,57^ (**Fig. 4a,c,d**). In addition, many transcripts encoding for molecules involved in cell survival and preventing apoptosis like BIRC3, VIM, and BCL2L1^58^ were also increased in cells from patients with severe COVID-19 illness, although some molecules with pro-apoptotic function (*e.g*., BAX, BIM, FAS)^59^ were also increased (**Fig. 4a-d** and **Extended Data Fig. 4b**). TCR sequence analysis indicated greater clonal expansion of cells in cluster 0 and 2 from severe COVID-19 patients (**Fig. 4e**). Overall, our findings suggest that SARS-CoV-2-reactive CD8_+_ T cells from patients with severe COVID-19 displayed multiple features that support the generation of robust CD8_+_ memory T cell responses with pro-survival properties.

**Figure 4.**
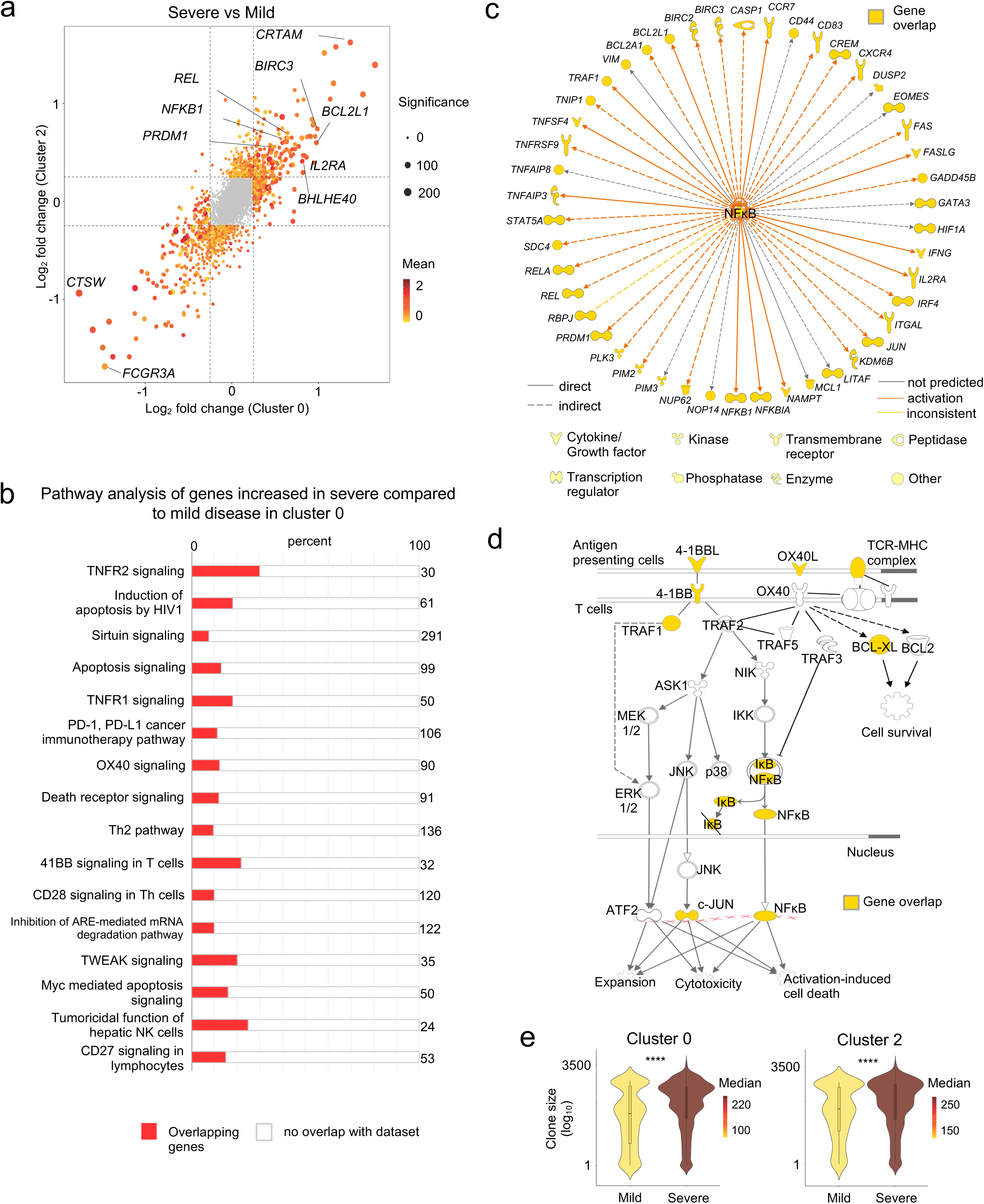
Pro-survival features in SARS-CoV-2-reactive CD8_+_ T cells from patients with severe COVID-19 illness. **a**, Plot shows fold change values of differentially expressed genes (adjusted *P* < 0.05, mean CPM >0, log_2_ fold change >0.25) in cluster 0 (x-axis) and 2 (y-axis) when comparing COVID-19 patients with severe and mild illness. A positive value indicates that the particular gene has increased expression in patients with severe disease relative to patients with mild disease in a given cluster, while a negative value indicates the opposite. **b-d**, Ingenuity pathway analysis (IPA) of genes with increased expression (adjusted *P* <0.05, log_2_ fold change >0.25) in cluster 0 cells between COVID-19 patients with severe versus mild illness; **b**, Top 16 canonical pathways with significant enrichment; **c**, Upstream regulatory network analysis of genes in NF-κB pathway; **d**, Transcripts encoding components in the 4-1BB and OX40 signaling pathway. **e**, Violin plots showing the degree of CD8_+_ T cell-clonal expansion in cluster 0 and 2 between COVID-19 patients with mild and severe disease. **** *P* <0.0001 by Mann-Whitney test.

## DISCUSSION

Recent studies in COVID-19 patients have verified the presence of CD8_+_ T cells that are reactive to SARS-CoV-2^5,7,8^. However, the nature and types of CD8_+_ T cell subsets that respond to SARS-CoV-2 and whether they play an essential role in driving protective or pathogenic immune responses remain elusive. Here, we report on the single-cell transcriptomes and TCR sequence analyses of >87,000 virus-reactive CD8_+_ T cells from 39 COVID-19 patients and 10 healthy, pre-pandemic donors. To compare the molecular properties of antigen-specific SARS-CoV-2-reactive CD8_+_ T cells to other common respiratory virus-reactive CD8_+_ T cells, we also isolated virus-reactive CD8_+_ T cells from healthy control subjects and analyzed their single-cell transcriptomes.

Across all the virus-reactive CD8_+_ T cells studied, we delineated eight distinct clusters with distinct transcriptomic features that shared TCRs, suggesting a high degree of plasticity among virus-reactive CD8_+_ T cells. We find that CD8_+_ memory T cells can acquire a wide range of transcriptional programs following different viral infections. For example, in healthy subjects, CD8_+_ T cells with polyfunctional features, linked to protective anti-viral immunity^36–39^, are abundant among CD8_+_ memory T cells reactive to FLU and RSV. In contrast, these cells were mostly absent in SARS-CoV-2 responsive cells from both COVID-19 patients and healthy non-exposed subjects. Notably, cells in this polyfunctional cluster were also significantly enriched for genes related to aerobic glycolysis, which is considered to enhance effector functions of CD8_+_ memory T cells^40,41^.

A large fraction of SARS-CoV-2-reactive cells (43% and 37%) from healthy non-exposed subjects (pre-pandemic), presumed to be human coronaviridae (CoV)-reactive cells that cross-react with SARS-CoV-2 peptide pools^5,21^, were present in clusters 1 and 0, respectively. These clusters also had similar representation of SARS-CoV-2-reactive cells from patients with COVID-19 illness. Cells in cluster 1 showed significant positive enrichment for type 1 interferon and ‘exhaustion’ signatures, reminiscent of the ‘exhausted’ CD8_+_ T cells reported in murine LCMV infection models^60^. Cluster 0, in contrast, was non-exhausted and showed significant negative enrichment of exhaustion and interferon signatures. SARS-CoV-2-reactive cells from COVID-19 patients also contributed the large majority of cells in cluster 2, which was characterized by enriched expression of cell cycle-related genes. Similar to cells in cluster 0, cluster 2 cells also showed negative enrichment of interferon genes and relatively lower exhaustion signature scores. Thus, we find that the nature of CD8_+_ T cells reactive to coronaviridae differed substantially from those responding to FLU or RSV.

Intriguingly, COVID-19 patients broadly segregated into two groups, according to whether the majority of the virus-reactive CD8_+_ memory T cells were in the ‘non-exhausted’ cluster 0 or the ‘exhausted’ cluster 1. Patients with mild COVID-19 illness had a greater proportion of SARS-CoV-2-reactive cells in the ‘exhausted’ cluster 1. Besides, cells in the ‘exhausted’ subset (cluster 1) from patients with mild COVID-19 illness expressed significantly lesser levels of transcripts encoding for cytotoxicity molecules, Fas ligand and pro-inflammatory cytokines (CCL3, CCL4, CSF-2, TNF, LTA, and LTB)^5,9,43,44^, and were significantly less clonally expanded. This finding raises the possibility that the magnitude and quality of the ‘exhausted’ CD8_+_ T cell response may be clinically important for limiting excessive tissue damage by SARS-CoV-2-reactive CD8_+_ T cells in COVID-19 illness.

Qualitative differences also emerged between patients with mild and severe COVID-19 illness in the ‘non-exhausted’ clusters, cluster 0 and 2. Transcripts increased in cluster 0 cells from patients with severe illness were significantly enriched in multiple co-stimulation pathways (OX40, CD27, CD28, 4-1BB, CD40), NF-κB, and cell survival pathways thought to be important for IL-2 production, proliferation, and survival. This finding suggested that patients with severe disease mount a more effective CD8_+_ memory T cell response to SARS-CoV-2 infection that could potentially lead to durable protection against re-exposure. Overall, our findings indicate that SARS-CoV-2-reactive CD8_+_ T cells from patients with severe COVID-19 displayed pro-survival properties and multiple features that potentially support the generation of robust CD8_+_ memory T cell responses and a lack of restraint reflected in the absence of an ‘exhaustion’ program. Whether these cells play a role in disease pathogenesis or provide long-term protection is not clear, and further longitudinal analysis and functional studies in relevant model organisms are required to clarify this.

## ONLINE METHODS

### Patient recruitment, ethics approval, and sample processing

The Ethics Committee of La Jolla Institute (USA) and the Berkshire Research Ethics Committee (UK) 20/SC/0155 provided ethical approval for this study with written consent from all participants. 22 hospitalized patients with reverse transcriptase polymerase chain reaction (RT-PCR) assay confirmed SARS-CoV-2 infection were recruited between April-May 2020. A cohort of 17 non-hospitalized participants was also recruited with RT-PCR assay or serological evidence of SARS-CoV-2 infection. Up to 80 ml of blood was obtained from all subjects for this research. Clinical metadata linked to hospitalized patients such as age, gender, comorbidities, level of clinical care required, radiological findings, and laboratory results are provided in **Extended Data Table 1**. The COVID-19 cohort consisted of 30 (77%) White British/White Other, 4 (10%) Indian, 2 (5%) Black British, 2 Arab (5%), and 1 Chinese (3%) participants. Of the 39 COVID-19 subjects, 22 (56%) had moderate/severe disease requiring hospitalization, and 17 (44%) had mild disease, not requiring hospitalization. The median age of hospitalized patients was 60 (33-82), and 77% were male. The median age of the non-hospitalized participants was 39 (22-50), and 47% were male. To study SARS-CoV-2-, FLU- and RSV-reactive CD8_+_ T cells from healthy subjects, we utilized de-identified buffy coat samples from 10 healthy adult subjects who donated blood at the San Diego Blood Bank before 2019, prior to the COVID-19 pandemic. Peripheral blood mononuclear cells (PBMC) were isolated from blood by density centrifugation over Lymphoprep (Axis-Shield PoC AS, Oslo, Norway) and cryopreserved in 50% human serum, 40% complete RMPI 1640 medium and 10% DMSO.

### Peptide pools

Two overlapping 15-mer SARS-CoV-2 peptide pools were purchased from Miltenyi Biotec^8^. These peptide pools were derived from the immunodominant sequence of the spike glycoprotein (S) and the complete sequence of the membrane glycoprotein (M) of SARS-CoV-2. For capturing FLU- and RSV-reactive CD8_+_ T cells, PepTivator Influenza A (H1N1) and RSV strain B1 peptide pools that covered the entire sequence of Hemagglutinin (HA) and Nucleoprotein (N) of each virus respectively, were obtained from Miltenyi Biotec.

### Antigen-reactive T cell enrichment (ARTE) assay

Virus-reactive CD8_+_ memory T cells were isolated using the protocol from Bacher *et al*. 2016 with minor modifications^15^. Thawed PBMC were plated overnight (5 % CO_2_, 37 °C) in 1 ml (concentration of 5 x 10^6^ cells/ml) of TexMACS medium (Miltenyi Biotec) on 24-well culture plates. Each of the SARS-CoV-2-specific peptide pools (1 μg/ml) were added separately to the PBMC culture for 24 hours. For subsequent magnetic-based enrichment of CD137_+_ cells, cells were sequentially stained with human serum IgG (Sigma Aldrich) for FcR block, cell viability dye (eFluor780/APC.Cy7, eBioscience), Fluorescence-conjugated antibodies, Cell-hashtag TotalSeq™-C antibody (0.5 μg/condition, clone: LNH-94;2M2, Biolegend), and a biotin-conjugated CD137 antibody (clone REA765; Miltenyi Biotec) followed by anti-biotin microbeads (Miltenyi Biotec). The following fluorescence-conjugated antibodies were used: anti-human CD3 (UCHT1, Biolegend), CD4 (OKT4, Biolegend), CD8B (SIDI8BEE, eBioscience), CD137 (4B4-1, Biolegend), CD69 (FN50, Biolegend), CCR7 (3D12, BD Biosciences), CD45RA (HI100, Biolegend), CD38 (HB-7, Biolegend), HLA-DR (G46-6, BD Biosciences), PD-1 (EH12.1, Biolegend). Antibody-tagged cells were added to MS columns (Miltenyi Biotec) to positively select CD137_+_ cells. After elution, FACSAria Fusion Cell Sorter (Becton Dickinson) was utilized to sort CD8_+_ memory T cells expressing CD137 and CD69. **Extended Data Fig. 1a** shows the gating strategy used for sorting. FlowJo software (v10.6.0) was employed for all Flow Cytometry Data analysis. In parallel, virus-reactive CD4_+_ memory T cells were isolated from the same cultures and analysis of their single-cell transcriptomes reported elsewhere^61^.

### Cell isolation and single-cell RNA-seq assay (10x platform)

To facilitate the integration of single-cell RNA-seq and TCR-seq profiling from the sorted CD8_+_ T cells, 10x Genomics 5’TAG v1.0 chemistry was utilized. A maximum of 60,000 virus-reactive CD8_+_ memory T cells from up to 8 donors was sorted into a low retention 1.5mL collection tube containing 500 μL of a solution of PBS:FBS (1:1) supplemented with RNAse inhibitor (1:100). After sorting, icecold PBS was added to make up to a volume of 1400 μl. Cells were then spun down (5 min, 600 g, 4 °C), and the supernatant was carefully aspirated, leaving 5 to 10 μl. The cell pellet was gently resuspended in 25 μl of resuspension buffer (0.22 μm filtered ice-cold PBS supplemented with ultra-pure bovine serum albumin; 0.04 %, Sigma-Aldrich). Following that, 33 μl of the cell suspension was transferred to a PCR-tube and single-cell library prepared as per the manufacturer’s instructions (10x Genomics).

### Single-cell transcriptome analysis

Using 10x Genomics’ Cell Ranger software (v3.1.0) and the GRCh37 reference (v3.0.0) genome, reads from single-cell RNA-seq were aligned and collapsed into Unique Molecular Identifiers (UMI) counts. The Feature Barcoding Analysis pipeline from Cell Ranger was used to generate hashtag UMI counts for each TotalSeq™-C antibody-capture library. UMI counts of cell barcodes were first obtained from the raw data output, and barcodes with less than 100 UMI for the most abundant hashtag were filtered out. Donor identities were assigned using Seurat’s (v3.1.5)^62^ *MULTIseqDemux* (autoThresh = TRUE and maxiter = 10) with the UMI counts. Cell barcodes were classified into three categories: donor ID (singlet), Doublet, Negative enrichment. Singlet cells were then stringently reclassified as doublet if the ratio of UMI counts between the top 2 barcodes was less than 3. All cells that were not classified as doublets or negative were used for downstream analysis. Cells from two COVID-19 patients with mild disease (patient 28 and 48) were not identifiable in the downstream analyses due to the lack of cell hashtags.

Single-cell RNA-Seq libraries (N = 15) were aggregated using Cell Ranger’s *aggr* function (v3.1.0). Analysis of the combined data was carried out in the R statistical environment using the package Seurat (v3.1.5). To filter out doublets and to eliminate cells with low-quality transcriptomes, cells were excluded if they were expressing < 800 or > 4400 unique genes, had < 1500 or > 20,000 total UMI content, and > 10% of mitochondrial UMIs. The summary statistics for each single-cell transcriptome library are given in **Extended Data Table 3** and show good quality data with no major differences in quality control metrics between batches (**Extended Data Fig. 1c**). Only transcripts expressed in at least 0.1% of the cells were included for further analysis. Using default settings in Seurat software, the filtered transcriptome data was then normalized (by a factor of 10,000) and log-transformed per cell. The top variable genes with a mean expression greater than 0.01 counts per million (CPM) and explaining 25% of the total variance were selected using the Variance Stabilizing Transformation method^62^. The transcriptomic data was then further scaled by regressing the number of UMIs detected, and the percentage of mitochondrial counts per cell. Principal component analysis was performed using the top variable genes, and based on the standard deviation of the principal components portrayed as an “elbow plot”, the first 25 principal components (PCs) were selected for downstream analyses. Cells were clustered using the *FindNeighbors*, and *FindClusters* functions in Seurat with a resolution of 0.2. The robustness of clustering was verified by other clustering methods and by modifying the number of PCs and variable genes utilized for clustering. Analysis of clustering patterns of SARS-CoV-2-reactive CD8_+_ T cells across multiple batches revealed no evidence of strong batch effects (**Extended Data Fig. 1d**). Plots to visualize normalized UMI data were created using the Seurat package and custom R scripts.

### Single-cell differential gene expression analysis

MAST package in R (v1.8.2)^63^ was used to perform pair-wise single-cell differential gene expression analysis after converting UMI data to log_2_(CPM+1). For genes to be considered differentially expressed, the following thresholds were used: Benjamini-Hochberg–adjusted *P*-value < 0.05 and a log_2_ fold change greater than 0.25. Cluster markers (transcripts enriched in a given cluster) were determined using the function *FindAllMarkers* from Seurat.

### Gene Set Enrichment Analysis and Signature Module Scores

Signature lists were extracted from published data sets and databases. Gene names from murine datasets were converted to human gene names using the biomaRt R package. Gene lists were then filtered to exclude genes that were expressed (CPM > 0) in < 2 % of the cells. Exhaustion consensus signature list was derived by considering genes that were present in > 3 exhaustion signature datasets^23–31^ (**Extended Data Table 5**). Genes that were present in cytotoxicity signatures^23,64^ or viral activation signatures^57^ were then excluded from this consensus list. The R package *fgsea* was used to calculate the GSEA scores with the signal-to-noise ratio as a metric. Default parameters other than minSize = 3 and maxSize = 500 were used. Normalized enrichment scores were presented as enrichment plots.

Signature scores were estimated with the Seurat’s *AddModuleScore* function, using default settings. For a given cell, each signature score is defined by the mean of the gene list of interest minus the mean expression of control gene lists. Control gene lists were sampled (size equal to the signature list) from bins defined based on the level of expression of the genes in the signature list. Signature gene lists used for analysis are provided in **Extended Data Table 5**.

### Single-cell trajectory analysis

Monocle 3 (v0.2.1)^65^ was used to calculate the “branched” trajectory; settings included the number of UMI and percentage of mitochondrial UMI as the model formula, and taking the highly variable genes from Seurat for consistency. After using the PCA output from Seurat and allocating a single partition for all cells, the cell-trajectory was outlined on the UMAP generated from Seurat as well. The ‘root’ was assigned using the *get_earliest_principal_node* function given in the package’s tutorial.

### T cell receptor (TCR) sequence analysis

Single-cell libraries enriched for V(D)J TCR sequences were processed to get clonotype information for each independent sample with the *vdj* pipeline from Cell Ranger (v3.1.0 and human annotations reference GRCh38, v3.1.0, as recommended). Joint analysis of single-cell transcriptomes and TCR repertoires was performed by aggregating independent libraries through custom scripts. For this purpose, cell barcodes were matched between corresponding libraries from each type. Then, each unique clonotype, a set of productive Complementarity-Determining Region 3 (CDR3) sequences as defined by 10x Genomics, was identified across all library annotations files. Finally, clone statistics, mainly clonotypes’ frequencies, and proportions were recalculated for the whole aggregation (considering only cells found in both modalities) so that previously-identified good quality cells were annotated with a specific clonotype ID and such clone statistics. Clone size was calculated as the number of cells expressing a given clonotype ID, and a clonotype was called as clonally expanded if this value was greater than 1 (clone size ≥ 2). Clone size was depicted on UMAP (per cell) or violin plots (per group, where color indicated clone size median of each group) using custom scripts, and clonotype sharing was presented using the tools from UpSetR^66^.

### Ingenuity Pathway Analysis (IPA)

IPA was performed using the default setting (v01-16) on transcripts that were significantly increased in expression in cluster 0 cells from patients with severe COVID-19 illness compared to mild illness. The canonical pathway analysis was performed to elucidate the enriched pathways in this data set and to visualize and highlight the gene overlap between the given data set with a particular enriched pathway. The upstream regulator network analysis was used to identify and visualize the interactions between differentially expressed downstream target genes in a given dataset with a particular upstream regulator.

### Statistical Analysis

Graphpad Prism 8.4.3 software was utilized for relevant data statistical analysis. Detailed information regarding statistical analysis, including test types and number of batches or samples is provided in the figure legends. P values are specified in the text or the figure legends. The data normality tests were performed, and for data that fell within Gaussian distribution, appropriate parametric statistical tests were performed. For those that did not conform to the equal variance-Gaussian distribution, appropriate non-parametric statistical tests were used.

### Data Availability

Scripts to obtain figures and tables provided in the manuscript are available in our repository on GitHub (https://github.com/vijaybioinfo/COVID19_2020). Sequencing data for this study has been deposited into the Gene Expression Omnibus under GSE153931.

## ACKNOWLEDGMENTS

We thank Luke Smith for patient recruitment and sample collection; Callum Dixon, Benjamin Johnson, Lydia Scarlett, and Silvia Austin for collection of clinical data; Céline Galloway, Oliver Wood, Katy McCann and Lindsey Chudley for sample processing. We thank the La Jolla Institute (LJI) Flow Cytometry Core for assisting with cell sorting; the LJI’s Clinical Studies Core for organizing sample collection. We thank Alessandro Sette for provided essential reagents for optimizing isolation of viral-reactive CD8_+_ T cells. Peter Friedmann and Anusha Preethi Ganesan for providing critical feedback on the manuscript. This work was funded by NIH grants U19AI142742, U19AI142742-02S1 (P.V., C.H.O), U19AI118626 (P.V., G.S.), R01HL114093 (P.V., F.A., G.S.,), R35-GM128938 (F.A), S10RR027366 (BD FACSAria-II), S10OD025052 (Illumina Novaseq6000), the William K. Bowes Jr Foundation (P.V.), and Whittaker foundation (P.V., C.H.O.). Supported by the Wessex Clinical Research Network and National Institute of Health Research UK.

## AUTHOR CONTRIBUTIONS

A.K., S.J.C., C.H.O., and P.V. conceived the work. A.K., S.J.C, C.H.O., and P.V. designed the study and wrote the manuscript. S.J.C supervised patient identification, recruitment, sample collection and processing. E.P., supervised the analysis of viral PCR and serology tests. A.K. and B.M., performed ARTE assay and FACS sorting, and H.S., performed single-cell RNA-sequencing under the supervision of C.H.O., and P.V. C.R.S. and V.F.R, performed bioinformatic analyses under the supervision of G.S., F.A., C.H.O., and P.V.

## Extended Data Figure Legends

**Extended Data Figure 1.**
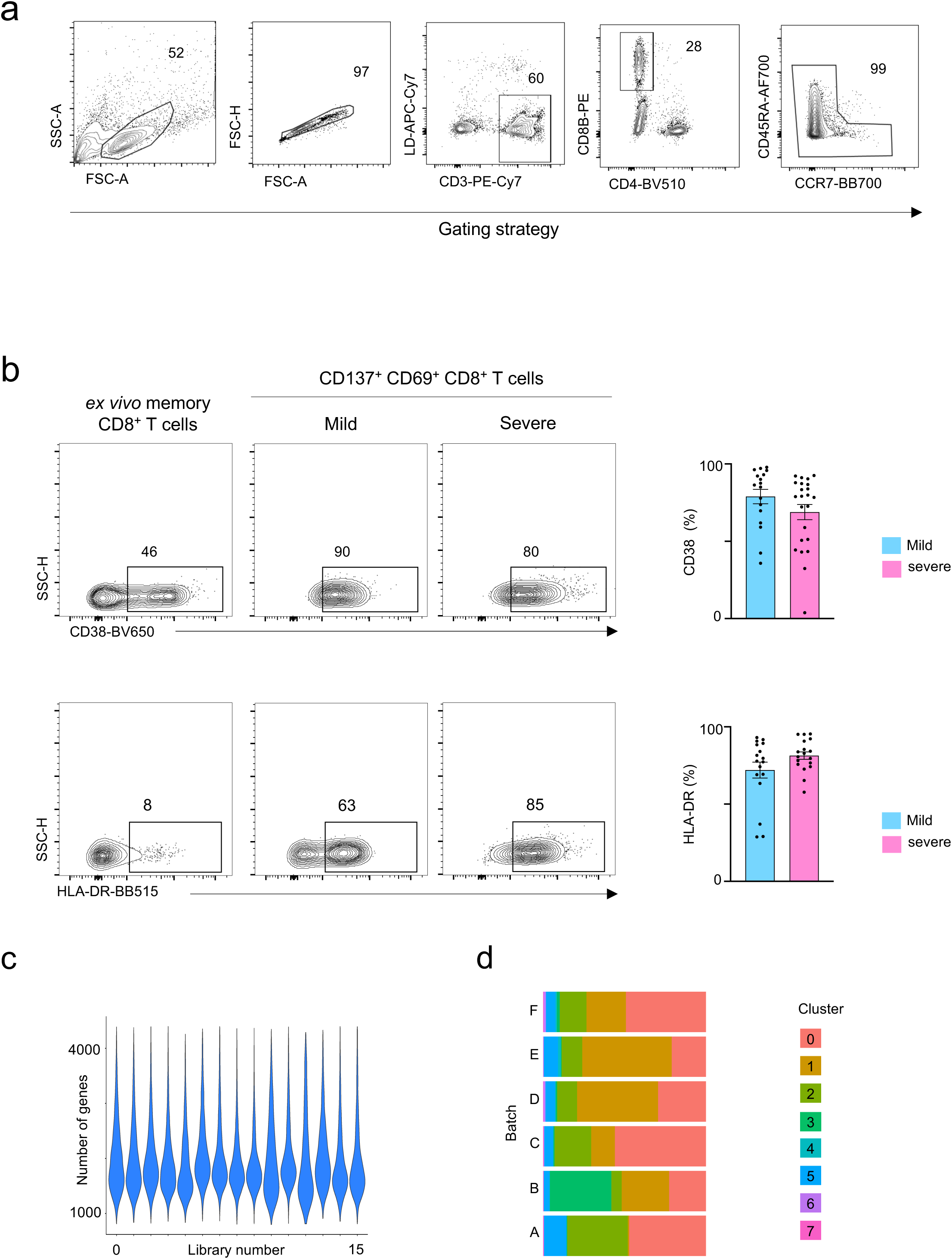
CD8_+_ T cell responses in COVID-19 illness. **a**, Gating strategy to sort: lymphocytes size-scatter gate, single cells (Height vs Area forward scatter (FSC)), live, CD3_+_CD8_+_ memory (CD45RA_+_CCR7_+_ naïve cells excluded) activated CD137_+_ CD69_+_ cells. Surface expression of activation markers was analyzed on CD8_+_ memory T cells. **b**, Representative FACS plots (left) showing surface expression of CD38 and HLA-DR in CD8_+_ memory T cells *ex vivo* and in CD137_+_CD69_+_ CD8_+_ memory T cells following 24 hours of stimulation, post-enrichment (CD137-based), and summary of proportions of cells expressing CD38 (N=39, P=0.24 by Mann-Whitney test)) and HLA-DR (N=34, P=0.11 by Mann-Whitney test) in CD137_+_CD69_+_ CD8_+_ memory T cells following stimulation and post-enrichment (CD137-based) in COVID-19 patients with mild and severe disease (right); Data are mean +/- S.E.M. **c**, Number of genes recovered for each 10X library sequenced. **d**, Distribution of cells in each cluster for the 6 batches of SARS-CoV2-reactive CD8_+_ T cells from COVID-19 patients (right panel).

**Extended Data Figure 2.**
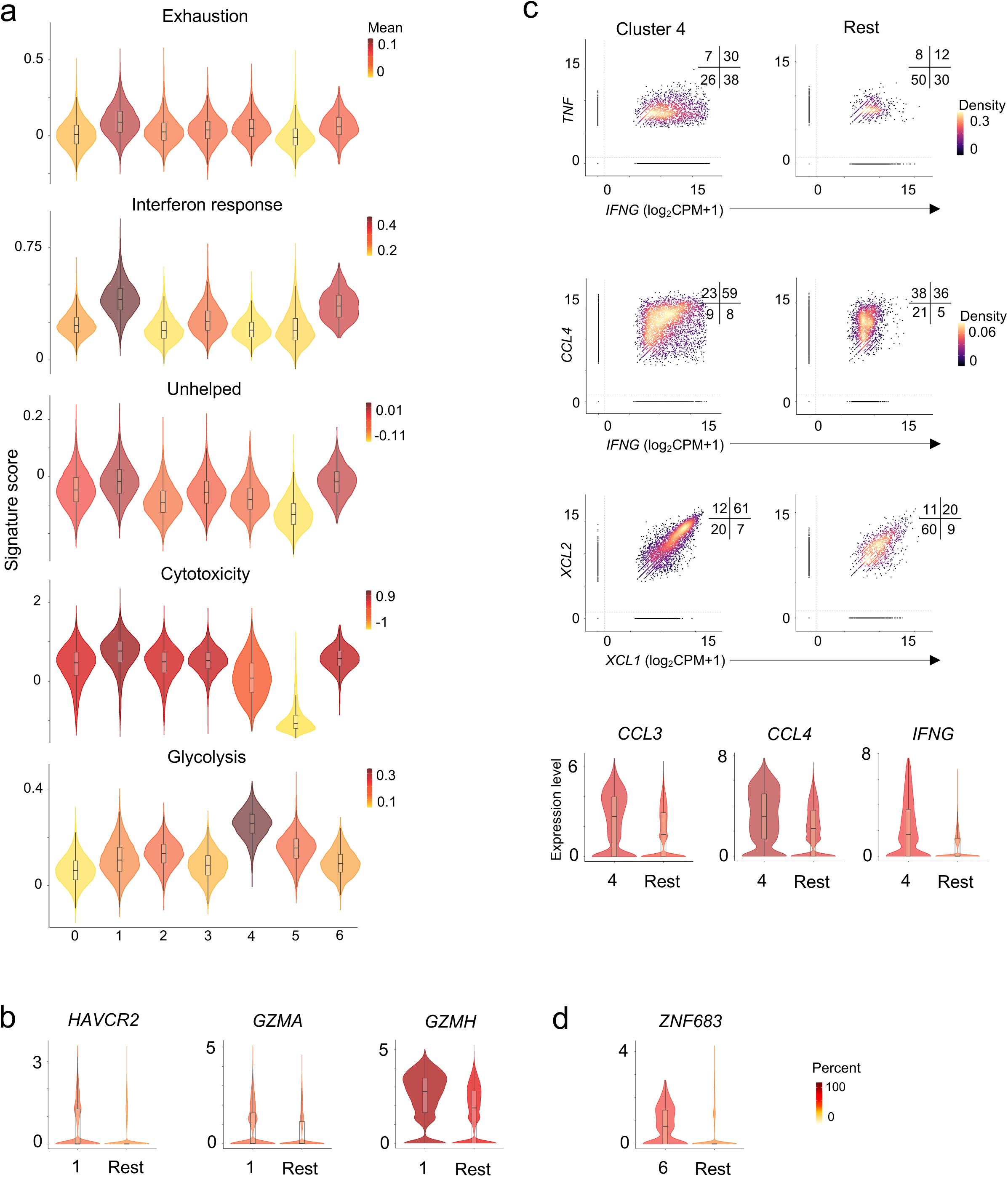
Virus-reactive CD8_+_ T cells show transcriptomic heterogeneity. **a**, Violin plots showing enrichment patterns of exhaustion, interferon (IFN) response, cytotoxicity, ‘unhelped’, and glycolysis gene signatures for each cluster. Color indicates signature scores. **b**, Violin plots showing expression of exhaustion and cytotoxicity gene markers in cluster 1 compared to an aggregation of remaining cells (Rest). **c**, Scatter plots (top panel) displaying co-expression of *IFNG* and *TNF, IFNG* and *CCL4, XCL1* and *XCL2* transcripts in virus-reactive CD8_+_ memory T cells in cluster 4 compared to the rest of the cells (Rest). Numbers indicate the percentage of cells in each quadrant. Violin plots (bottom panel) showing expression of indicated transcripts in cluster 4 compared to an aggregation of remaining cells (Rest). **d**, Violin plot showing expression of *ZNF683* in cluster 6 compared to an aggregation of remaining cells (Rest). Color indicates percentage of cells expressing indicated transcript.

**Extended Data Figure 3.**
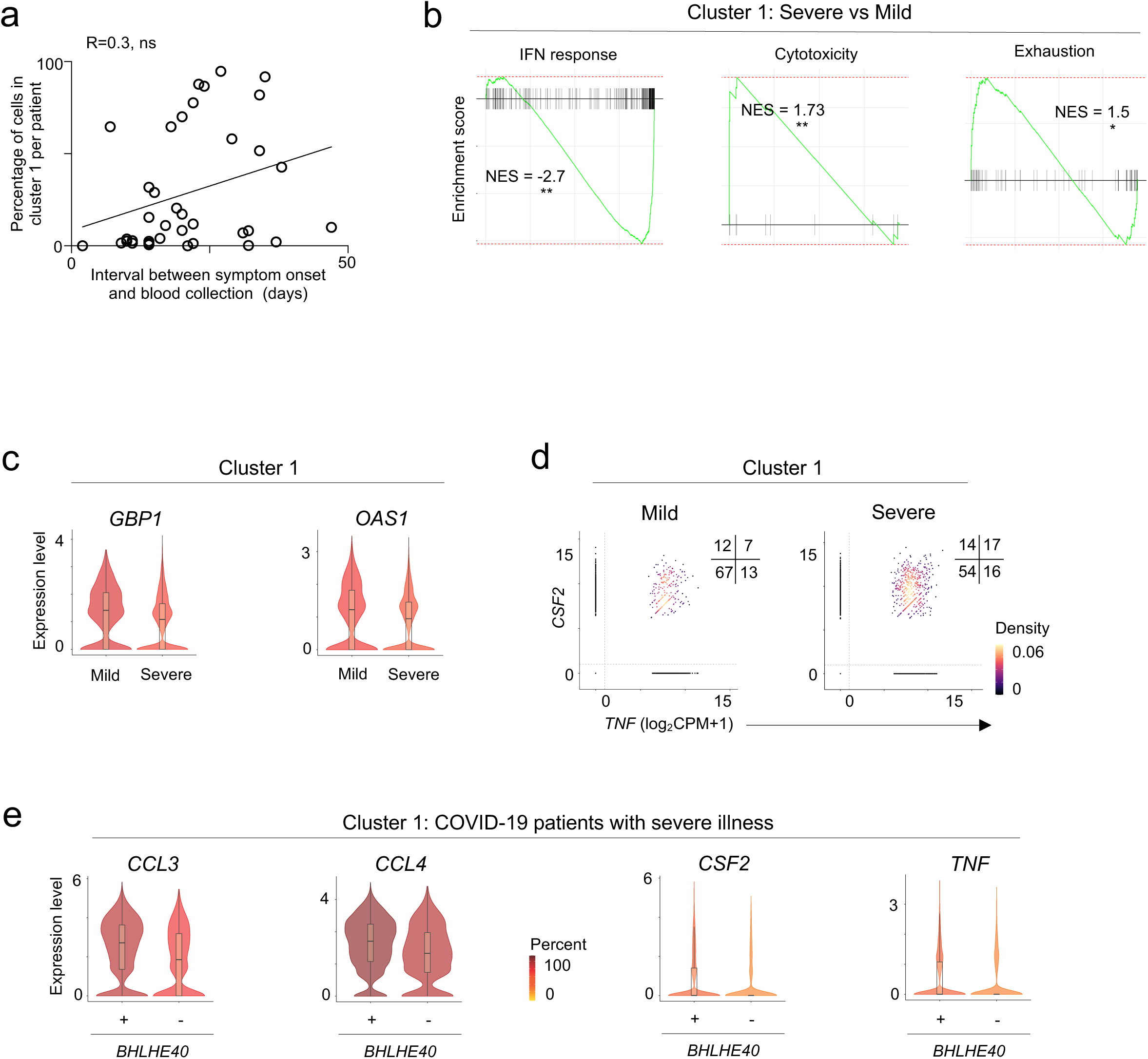
Exhausted SARS-CoV-2-reactive CD8_+_ cells are increased in mild COVID-19 illness. **a**, Plot depicting correlation of the proportion of cells in cluster 1 per COVID-19 patients (y-axis, percent) and the interval between symptom onset to blood collection (x-axis, days). R=0.3 (Pearson correlation), P<0.07 (ns). **b**, Gene Set Enrichment Analysis (GSEA) of Interferon response, Cytotoxicity, and Exhaustion signatures in cluster 1 cells between COVID-19 patients with severe versus mild illness. **c**, Violin plots depicting several IFN response genes in cluster 1 cells from COVID-19 patients with mild and severe illness. **d**, Scatter plot displaying co-expression of *TNF* and *CSF2* transcripts in cluster 1 cells from COVID-19 patients with mild and severe illness. Numbers indicate the percentage of cells in each quadrant. **e**, Violin plots comparing the expression of indicated transcripts between *BHLHE40*-expressing and non-expressing cells in cluster 1 from COVID-19 patients with severe disease.

**Extended Data Figure 4.**
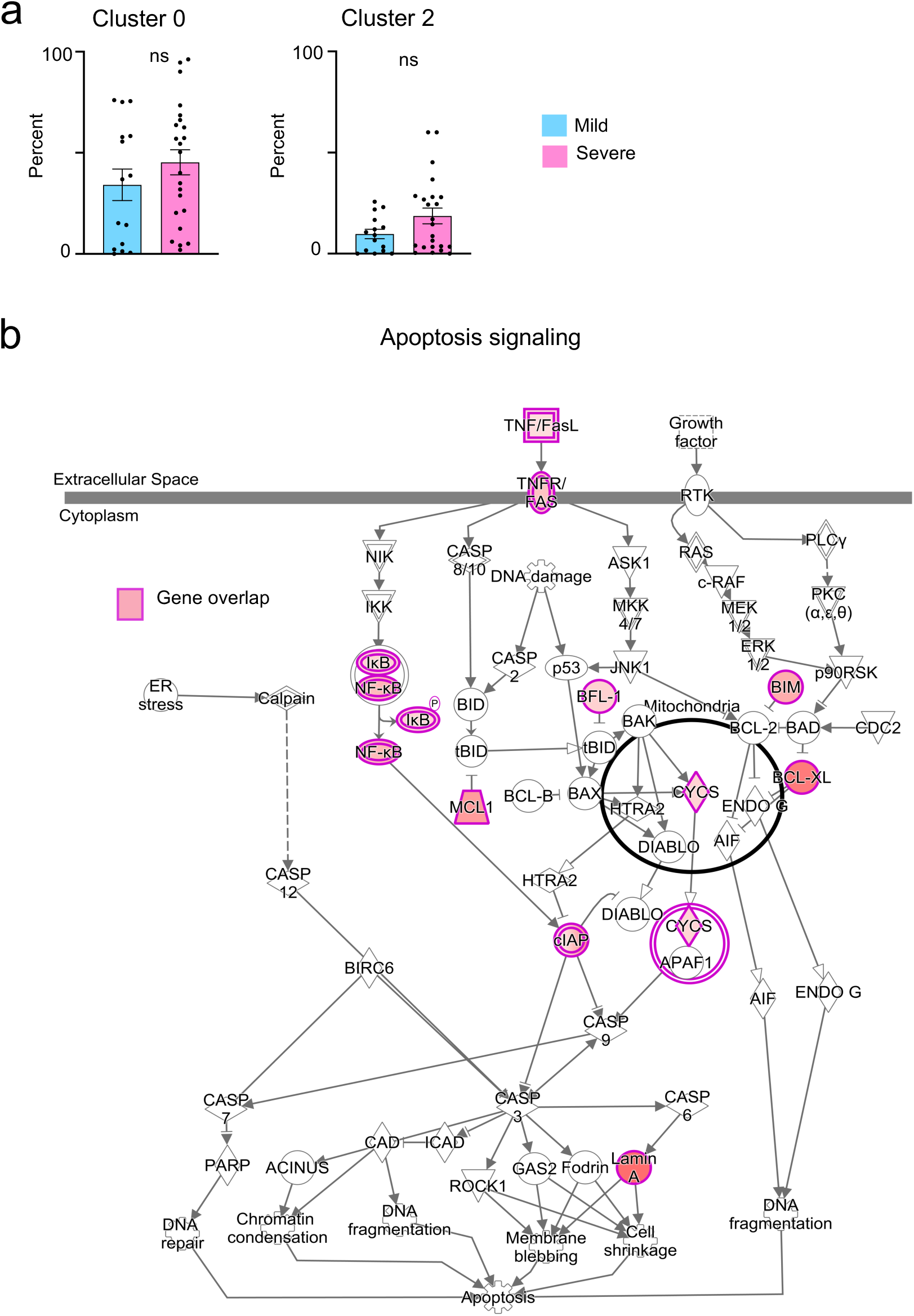
Pro-survival features in SARS-CoV-2-reactive CD8_+_ T cells from patients with severe COVID-19. **a**, Bar charts comparing the proportion of cells in cluster 0 (*P* = 0.29 by Mann-Whitney test) and 2 (*P* = 0.13 by Mann-Whitney test) from COVID-19 patients with mild and severe disease. Data are displayed as mean +/- S.E.M (N=37). **b**, Ingenuity pathway analysis (IPA) of genes with increased expression (adjusted *P* <0.05, log_2_ fold change >0.25) in cluster 0 cells between COVID-19 patients with severe versus mild illness; transcripts encoding components of apoptosis signaling pathway are shown.

## EXTENDED DATA TABLES

**Extended Data Table 1. Human subject details**

**Extended Data Table 2. Summary of all FACS data**

**Extended Data Table 3. Single-cell sequencing quality controls and subject-specific cell numbers**

**Extended Data Table 4. Single-cell cluster enriched transcripts**

**Extended Data Table 5. Gene lists utilized for Gene Set Enrichment Analysis and Signature Module Scores**

**Extended Data Table 6. Subject-specific TCR clonotype data**

**Extended Data Table 7. Cluster-specific TCR clonotype data**

**Extended Data Table 8. Single-cell differential gene expression analysis**

**Extended Data Table 9. Ingenuity Pathway Analysis**

